# Langevin Dynamics Simulation of AKAP-PKA Complex: Re-Envisioning the Local Concentration Mechanism for Directing PKA Phosphorylation

**DOI:** 10.1101/161760

**Authors:** Marc Rigatti, Paul J. Michalski, Kimberly L. Dodge-Kafka, Ion I. Moraru

## Abstract

The second messenger cAMP and its effector cAMP-dependent protein kinase A (PKA) constitute a ubiquitous cell signaling system. In its inactive state PKA is composed of two regulatory subunits that dimerize, and two catalytic subunits that are inhibited by the regulatory subunits. Activation of the catalytic subunits occurs upon binding of two molecules of cAMP to each regulatory subunit. Although many receptor types existing within the same cell may use this signaling system, compartmentation of signaling is thought to occur due to A-Kinase Anchoring Proteins (AKAPs), which act to co-localize PKA with specific substrates. However, the molecular mechanism allowing AKAPs to direct PKA phosphorylation to a particular substrate remained elusive, as prior evidence suggested that the catalytic subunit, which is highly diffusible, is released after cAMP binding to the regulatory subunit. Recent evidence from Smith et al. suggests that in the cell, the catalytic subunit may in fact not be released from the AKAP complex [1, 2]. They further demonstrated that alterations in the structure of the PKA regulatory subunit tether affect substrate phosphorylation. We use a novel computational software based on Langevin dynamics, SpringSaLaD, to simulate the AKAP-PKA complex in order to determine a molecular mechanism for the changes in phosphorylation seen with alteration in tether length and flexibility, and to demonstrate whether or not AKAPs can effectively direct PKA phosphorylation to a particular substrate upon release of the catalytic subunit from the complex. We find that short and flexible tethers contribute to a decrease in the average characteristic time of binding, allowing the catalytic subunit to spend more time in a bound state with the substrate, which yields faster characteristic times of phosphorylation. We further demonstrate that release of the catalytic subunit from the AKAP complex abrogates the effect of tethering, with characteristic times of phosphorylation similar to non-AKAP bound PKA. The data demonstrates that AKAPs likely do not release the catalytic subunit in directing PKA phosphorylation to AKAP bound substrates. In combination with the changes in characteristic time of phosphorylation which are driven by tether structure, this work indicates that the purpose of AKAPs may be to increase the efficiency of phosphorylation of particular AKAP substrates.

## Introduction

The second messenger cAMP and its effector, cAMP-dependent protein kinase A (PKA), constitute a ubiquitous cell-signaling network responsible for the phosphorylation of a vast array of substrates. Recent experimental methods employing mass spectrometry have identified an astounding 112 potential PKA substrates existing in a single cell type [3]. Such an abundance of PKA substrates in a single cell highlights the importance of signal compartmentation within the cAMP-PKA signaling pathways. We have known since the 1970’s that cAMP-PKA signaling is compartmentalized. Stimulation of the β-adrenergic receptor in cardiomyocytes was shown to the activate glycogen phosphorylase, while stimulation of the PGE1 receptor did not, even though both receptors signal via cAMP and PKA [4]. Historically, signal compartmentation was thought to arise from the multiplicity of PKA, and adenylate cyclases (ACs) and phosphodiesterases (PDEs) isoforms that produce and degrade cAMP respectively [5, 6]. Co-localization of the proper combination of AC, PKA, and PDE produces a spatially restricted cAMP signal that activates the local PKA pool resulting in phosphorylation of specific PKA substrates. While the notion of spatially restricted cAMP signals has been demonstrated and is well accepted, the structural basis for cAMP-PKA signal specificity is not well understood [7]. The most recent evolution of the compartmentation hypothesis involves assembly of a macromolecular complex or signal microdomain by A-Kinase Anchoring Proteins (AKAPs) [8]. Simultaneous binding of AKAPs to AC, PDE, PKA, and a specific substrate is thought confer spatiotemporal control of cAMP-PKA signals. Unfortunately, even this scheme does not lend a complete and satisfactory explanation of PKA signal specificity.

Achieving phosphorylation of specific substrates in response to an extracellular stimulus is complicated by the rapid diffusion of cAMP and also by diffusion of the PKA catalytic subunit. The vast majority of evidence regarding PKA activation indicates that binding of cAMP to allosteric sites on the regulatory subunits results in the simultaneous activation and release of the catalytic subunit [9]. The co-discoverer of cAMP, Ted Rall, once said that this scheme presents “…the unsatisfying picture of the catalytic subunit of protein kinase swimming about, happily phosphorylating a variety of cellular constituents whether they need it or not” [7]. Co-localization of PKA with its substrates according to the AKAP-PKA microdomain hypothesis is thought to confer specificity of phosphorylation by increasing the local concentration of PKA. While achieving a high local concentration of PKA via anchoring to a specific substrate in a particular location is straightforward, the result of this mechanism following PKA activation and release is less obvious. Since the activated catalytic subunit is considered to be freely diffusible, it may diffuse to and phosphorylate other substrates in the same manner expected for the AKAP bound substrate. While phosphorylation of non-local substrates may be inhibited by the presence of protein kinase inhibitor (PKI), a pseudosubstrate inhibitor of PKA, it must be acknowledged that PKA will not necessarily diffuse to and phosphorylate its associated substrate even if released in close proximity of its substrate. Thus, these data call into question the mechanism of PKA scaffolding.

We have taken a computational approach to investigate the effect of tethering PKA to its substrate, via formation of an AKAP-PKA microdomain, on signal specificity given the currently accepted mechanism of PKA activation and release. Using a novel software package employing Langevin Dynamics simulation, SpringSaLaD [10], we have demonstrated that co-localization of PKA and substrate within an AKAP complex does not reliably direct the catalytic subunit to substrates within the complex if the catalytic subunit is released. This suggests that to obtain the substrate specificity seen in multiple experimental models, PKA must retain its holoenzyme conformation.

Interestingly, recent experimental evidence has surfaced that PKA is not released following stimulation and is able to maintain catalytic activity while bound to its regulatory subunit [1, 2, 11]. The ability of PKA to phosphorylate its substrates while remaining bound to the AKAP complex would provide for a much more convincing hypothesis regarding specificity of substrate phosphorylation. This novel mechanism of PKA activation has recently been investigated by cryo-EM using AKAP7γ and AKAP5 as model systems [1, 2]. Additionaly, using proximity ligation assays between the Regulatiory and Catalytic subunit as well as immunoprecipitation and FRET experiments found no disruption of the PKA complex under physiological concentrations of cAMP [1]. The AKAP7γ-PKA complex has been demonstrated to posses a remarkable amount of conformational flexibility owing to the intrinsically disordered tether region of the PKA regulatory subunits [2]. This tether restricts the catalytic subunit to space within ~16nm of the AKAP-PKA interface. Truncation of the tether was shown to increase substrate phosphorylation relative to phosphorylation with the WT tether length. It was further demonstrated that following stimulation with the β-adrenergic agonist isoproterenol, the catalytic subunit remains within the AKAP complex. These results suggest that the tether physically constrains PKA to the location of the substrate during phosphorylation and that the structure of the tether is an important determinant of substrate phosphorylation.Importantly, these findings suggest that scaffolding of PKA to substrates by AKAP may not function as currently hypothesized.

In addition to demonstrating the ineffectiveness of co-localizing PKA and substrate given release of the catalytic subunit, we are able to provide a mechanistic explanation for the results of Smith *et al* by determining how tether length and flexibility are able to affect phosphorylation. Creation of a model of the AKAP7γ-PKA-substrate complex using SpringSaLaD allowed us to explore the effect modulating the length and flexibility of the PKA tether region on phosphorylation of AKAP complex bound substrate. Our work demonstrates that shorter and more flexible tethers allow the catalytic subunit to interact with its substrate more often, resulting in higher rates of phosphorylation. Importantly, the results show that PKA rarely phosphorylates its substrate within the first binding event, lending further support to our assertion that the catalytic subunit must remain bound within the complex.

Our findings strongly support emerging evidence that the PKA catalytic subunit is retained within the AKAP-PKA complex. Interestingly, the same mechanism that explains substrate specificity of the AKAP complex also seems to confer high catalytic efficiency, helping to explain the observation that AKAPs increase the speed of substrate phosphorylation [12]. This work lends new meaning to the idea of local concentration that better explains the ability of AKAPs to control substrate specificity.

## Methods

### Langevin Dynamics Modeling

LD simulations were carried out using *SpringSaLaD* [10], a novel open source software for mesosocopic simulations of molecular interactions; the latest version and documentation is available at http://www.ccam.uchc.edu/ccam-software/. *SpringSaLaD* is an appropriate modeling platform when the molecules in the system can be described in a coarse-grained manner as a set of biochemically distinct spherical sites connected by stiff links. The AKAP-PKA complex is well described in this manner. Models that seek to describe finer details, such as the motion of individual amino acids, are more appropriately simulated with molecular dynamics, while models with less detail will run faster with other solvers.

Molecules are constructed in the graphic user interface in a three-step process. First, the *types* of sites in the molecule are defined. For the AKAP-PKA complex, there are five types of sites: an anchor, which is necessary for linking molecules to a membrane, substrate, AKAP_R (AKAP bound to the regulatory subunit dimer), R_C (regulatory subunit bound to the catalytic subunit), and flex links that allow us to modulate flexibility of the regulatory subunit tether. Each type can possess an arbitrary number of internal states, which can be associated with a set of biochemical reactions (reactions are described below). For example, substrate is assigned two states, unphosphorylated and phosphorylated. Each type also has an associated physical size, diffusion constant, and color (for visualization purposes).

Secondly, *sites* are added to the molecule, and each site is assigned one of the previously defined types. To construct the AKAP-PKA complex we add a minimum of seven sites: an anchor, substrate, AKAP_R, two flex links, and two R_C. Physically, sites are modeled as impenetrable spheres in order to accurately capture excluded volume effects.

Lastly, *links* are added to connect the sites to each other. Each site can be linked to an arbitrary number of other sites in either two or three-dimensions. The only requirement is that sites cannot overlap. Links are stiff and thus define an inter-site distance, but are free to rotate around the sites. For example, a triangular molecule with three sites will not maintain its geometry in the simulation with only two links, but will if a third bond is added to enforce a distance between the two outer sites. Links are only used to control the distance between sites and do not occupy physical space, and both sites and other links are free to pass through a link. Excluded volume is only enforced by sites.

The AKAP-PKA complex model consisted of a single molecule composed of five types with a structure based on the published geometry of the complex as detailed in the results [2]. The governing equations and algorithms are discussed extensively in [10] and briefly described below. For each model variant, a 1sec simulation of a single complex in a 50nm^3^ volume was run 500 times on a high performance computing cluster. The phosphorylation state of the complex was recorded at 10ms intervals yielding 100 data points per simulation.

### Particle Motion: Diffusion and Constraints Due to Binding

Particle motion is influenced by two classes of forces: random forces that lead to diffusional motion, and inter-particle forces that impose the constraints from intra- and inter-molecular bonds. These forces are incorporated in the over-damped Langevin equation [1, 2],

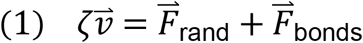

where ζ is the coefficient of viscous friction, 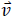 is the particle velocity, and 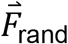 and 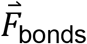 represent the random forces and the total force of bonds. The random forces are guaranteed to recapitulate the desired diffusion provided they are chosen from normal distribution with variance

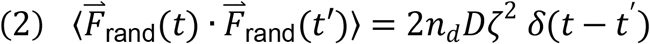

where D is the desired diffusion coefficient, *n* _*d*_ is the dimension of the system (here *n*_*d*_ = 3), and the delta function simply states that the random force is uncorrelated in time. The bonds are modeled as stiff springs,

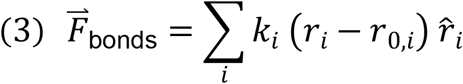

where the sum runs over all bonds, *k*_*i*_ is a spring constant, *ȓ*_*i*_ is the unit vector pointing from the particle to the neighbor with which it shares a bond, *r*_*i*_ is the current inter-particle distance, and *r*_0,*i*_ is the equilibrium bond distance. The exact value of the spring constant is not important, provided the spring is stiff enough to keep sites near the expected distances but not so stiff that accurate motion requires unreasonably small time steps. In SpringSaLaD all bonds have a spring constant that allows us to use time steps on the order of 1 to 100 ns for biologically relevant diffusion constants.

### Reactions

SpringSaLaD simulations support rate-controlled particle creation (zeroth order reactions) as well as first- and second-order reactions. Two general types of first order reactions can be specified in the model, both of which are described by a reaction rate *r*, with units s^-1^, and which occur at each time step with probability *rdt*. These are: 1) Bond dissociation reactions. When a dissociation reaction occurs the bond is simply removed from the system. 2) Internal state transitions. These describe the transitions between the internal states of each site, as defined by the type of that site. The probability of these transitions can depend on the identities of binding partners or the states of other sites in the same molecule. The former dependency would be used to prevent a transition from an unphosphorylated to a phosphorylated state unless the site is bound to a kinase. The latter dependency can be used to model allosteric interactions.

SpringSaLaD defines binding reactions between two sites as second order reactions. A bond is modeled by the creation of a new link between the reacting sites. The link modeled identically to the links which hold molecules together, except it has an associated off rate which controls molecular dissociation. Particle-based simulations often use the Smoluchowski approach [13] to model bimolecular reactions, but such an approach is incompatible with excluded volume. Instead, we modified the approach described in Ref. [14] to account for excluded volume. Each site is associated with two radii, the physical radius, ρ_*i*_ (i=1,2), which is defined in molecule construction and enforces excluded volume, and a slightly larger reaction radius, *R*_*i*_. Two reactive sites undergo a binding reaction with probability λ *dt* per time step when their reaction radii overlap. The reaction rate λ is related to the macroscopic on rate, *k*_on_, with units of μ*M*^−1^*s*^−1^, through

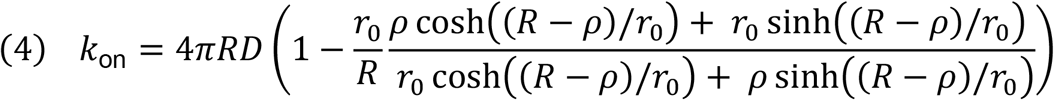

where 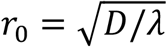, *D* = *D*_1_ + *D*_2_, ρ = ρ_1_ + ρ_2_, and *R* = *R*_1_ + *R*_2_.

### Data reduction and analysis

Data reduction and analysis was performed using custom scripts written in Python using the SciPy library. To determine the characteristic time of binding or phosphorylation, 500 runs of a single model were grouped into sets of 50 runs and the average for the unbound or unphosphorylated state was taken at each time point. Since there is only 1 molecule per simulation, this produces an exponential decay from 1 to 0 as the average state of the molecule transitions from completely unbound or unphosphorylated to completely bound or phosphorylated. The resulting 10 sets of averaged data produced from the 500 runs were then fit with a single exponential model (Eq. 5), where y is the average value of the state, b is the inverse of the characteristic time of binding or phosphorylation, and t is time.

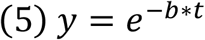

The mean and standard deviation of the characteristic times for the 10 sets of averaged data are reported.

To determine the average time spent in the bound state for a given set of simulations, the total number of data points in which the complex existed in a bound state was determined for each run. The time spent in the bound state was then averaged for all 500 simulations of a particular model. The mean and standard deviation are reported for each model variation.

### Analytical Prediction of Phosphorylation Rate

Expected catalytic rates for PKA were calculated using the standard Michaelis-Menten equation (Eq. 6), where V is reaction velocity (μm/s), k_cat_ is the catalytic rate (s^-1^), Km is the Michaelis constant ((μM), and [E] and [S] are the enzyme and substrate concentrations ((μM) respectively.

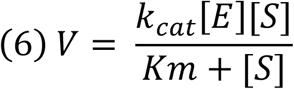

This model of ensemble enzymatic reactions has been shown, both theoretically and experimentally, to hold for single molecule reactions [15, 16]. The Michaelis constant was calculated from the following equation (Eq. 7), where kcat is the catalytic rate and kf and kr are the reverse and forward rate constants for the binding of enzyme to substrate.

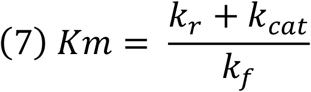

Since the volume accessible to the catalytic subunit is limited by the tether length (distance from center of the AKAP_R subunit to center of RC), the concentrations of E and S were adjusted to a spherical volume with a radius equal to the tether length. Given the adjusted concentrations and known k_cat_, k_f_ and k_r_, we determine V (μM/s). The reaction velocity is then converted to catalytic rate by dividing through by the adjusted enzyme concentration.

## Results

### Modeling the AKAP-PKA complex

Simulation of substrate phosphorylation within the AKAP-PKA complex is a multifaceted task that necessitates accounting for structural details in addition to standard reaction kinetics. Such structural detail generally falls within the realm of molecular dynamics simulation; however, this method is very computationally expensive and would not be appropriate given the time-scales of the binding and phosphorylation reactions in question. Similarly, stochastic simulation of this reaction network could be accomplished using a tool like Smoldyn. Yet, this method does not account for space filling of molecules and the spatial resolution of this type of modeling would be pushed to its absolute limits. Our novel simulation software takes advantage of Langevin Dynamics (LD) to provide a meso-scale modeling platform that accounts for coarse-grained structural details and is capable of simulating the AKAP-PKA complex with the appropriate spatial resolution.

In the LD simulations, molecules are constructed from spheres or *sites* of a particular type and springs or *links*. To demonstrate model construction with SpringSaLaD we will use our model of the AKAP7γ-PKA-substrate complex, which is constructed of 5 molecular *types*: anchor, substrate, AKAP_R, linker, and RC (Fig 1A). The geometry of the AKAP complex is based upon the 3D reconstruction from Smith *et al* [2]. In order to link the molecule to the membrane it is necessary to include an anchor, which represents the membrane insertion of our target substrate. The interface of AKAP with the dimerization/docking domain of the PKA regulatory subunits is represented by AKAP_R, and the catalytic subunit bound to the inhibitory domain of the regulatory subunit is represented by RC. The diffusion rates of the molecule types are based upon standard rates for cytosolic and membrane bound proteins. The diffusion rate of GFP in chinese hamster ovary cells has been measured at 27μm^2^/s [17]. Although our diffusion coefficient for RC is roughly one order of magnitude less than that of GFP, simulations with diffusion coefficients varying from 3-15μm^2^/s show no effect on the characteristic time of phosphorylation (data not shown). Diffusion rates for transmembrane proteins are generally less than 0.1μm^2^/s. Since links between sites are stiff, we create flexibility in the link between AKAP_R and RC by adding dummy sites called flex links. By increasing the number of links in the tether, we are able to increase the flexibility. To simulate the effects of tether length and flexibility on the rate of substrate phosphorylation, we created a series of models in which the total length of the tether (center of AKAP_R to center of RC) is varied as well as the number of link segments (Fig 4.1B). A total of 190 model variations were created with lengths varying by increments of 1nm from 9.25m to 25.25nm, and flexibility ranging from a minimum of 2 link segments to a maximum of 14 link segments. For shorter tether lengths, the maximum number of link segments is limited by the number of flex link sites that could be fit within the tether length. The properties assigned to each site type as well as the kinetics of the simulated reactions are outlined in Figure 4.1C. The reaction kinetics are based on electronic measurements of single-molecules catalysis by PKA (*18*).

**Figure 1.**
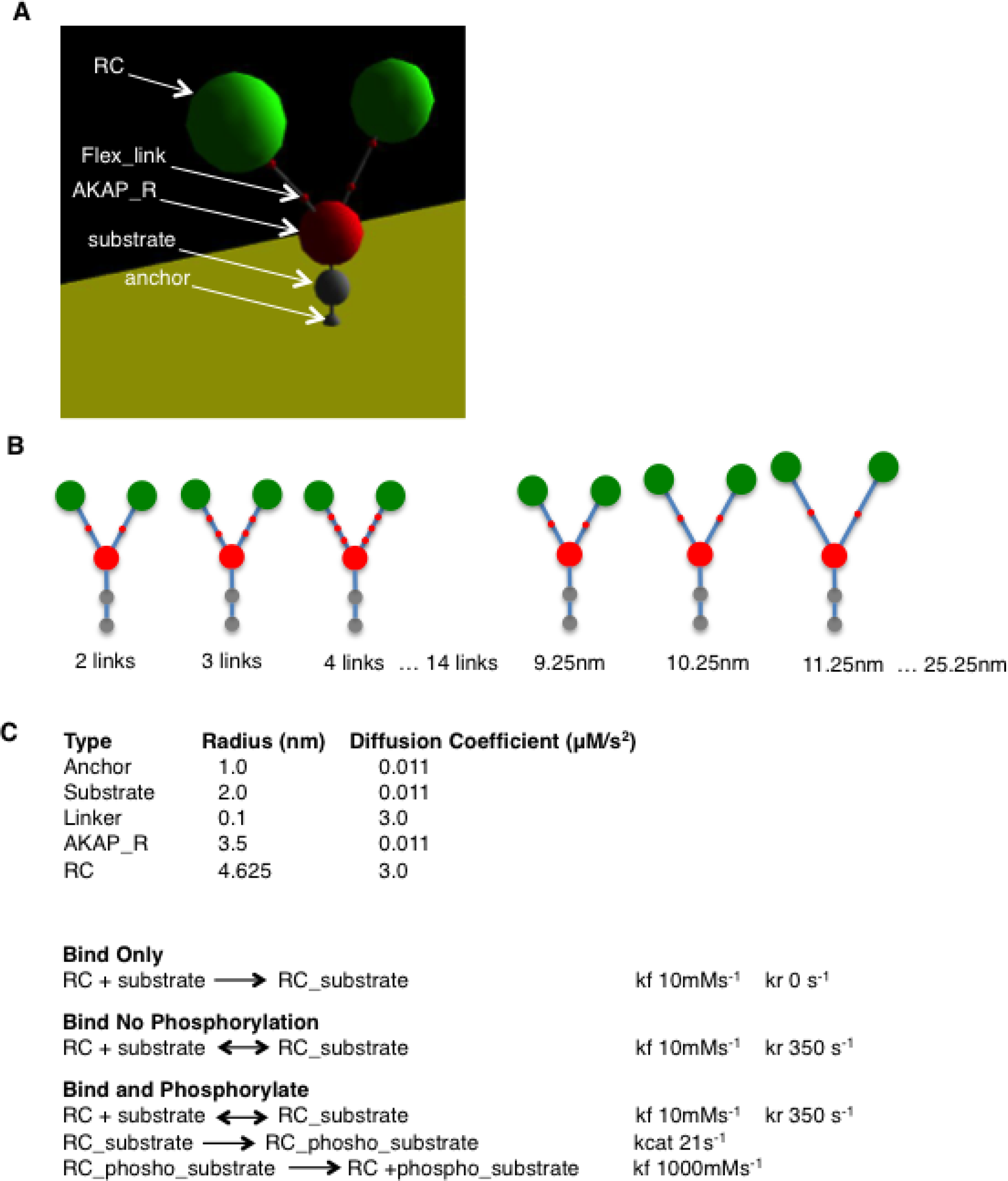
Langevin Dynamics Model of the AKAP-PKA complex. A) Structure of the AKAP-PKA complex with labeled site types. B) A total of 190 model variants were created that vary in both flexibility (left) and length (right). C) Model Parameters. Characteristics assigned to each site type (above) and kinetic parameters of each reaction network (below).

### Release of the catalytic subunit abrogates the effect of PKA tethering for a single AKAP-PKA-substrate complex

It is well established that the primary function of AKAPs is to compartmentalize cAMP-PKA signaling. Numerous studies have demonstrated that disruption of AKAP binding to a particular substrate leads to a decrease in phosphorylation or change in the expected physiologic response [18-20]. How AKAPs direct PKA phosphorylation to AKAP complex bound substrates remains unknown as the widely accepted mechanism of signaling requires the release of the catalytic subunit [9]. We hypothesized that given the free diffusion of the catalytic subunit upon release from the complex, efficient phosphorylation of AKAP complex bound substrates is not possible. Simulation of the release of the catalytic subunit from the complex allowed us to examine its effect on substrate phosphorylation.

To simulate the effect of releasing of the catalytic subunit on phosphorylation of AKAP complex bound substrate, we removed the tether from the AKAP7γ-PKA-substrate model described above, and initialized the simulations with two catalytic subunits positioned within 16.25nm of AKAP_R (WT), within 5nm of the substrate (close), or randomly throughout the 50nm^3^ simulation space (Fig 2A). Positioning of the catalytic subunits at the maximum distance from AKAP_R that could be expected for the wild type geometry lead to an average characteristic time of phosphorylation (CTP) of 0.193s ±0.025.

**Figure 2.**
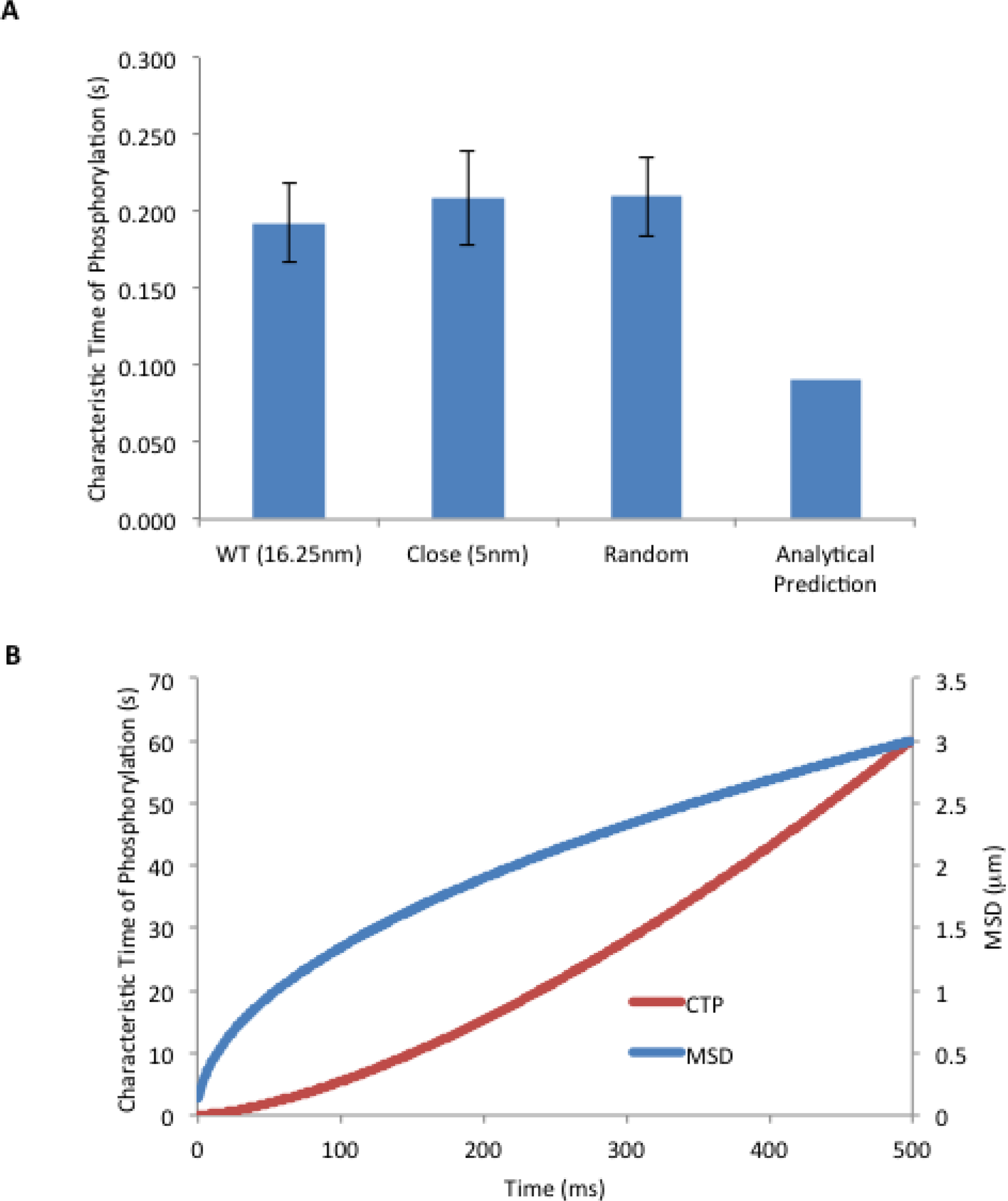
Simulation of release of the catalytic subunit from the AKAP-PKA complex. A) Release of the catalytic subunit was simulated by removing the tether from the model and initializing the simulations with the catalytic subunits positioned either 16.25nm from substrate, 5nm from substrate or randomly within the simulation space. The characteristic times of phosphorylation are compared to the analytical prediction for two catalytic subunits and a single substrate within a 50nm^3^ volume. B) Predicted change in characteristic time of phosphorylation (CTP) and mean squared distance of PKA as two catalytic subunits diffuse from their initial position within the AKAP complex.

Interestingly, positioning the catalytic subunits close enough to the substrate for binding to occur in the first time step of the simulation did not improve the CTP (0.208s ±0.031). Positioning the catalytic subunits within the space available to tethered PKA also did not improve the CTP compared to random positioning of the two catalytic subunits within the simulation space (0.210s ±0.026). These results suggest that anchoring of PKA to substrates by AKAP is not an effective method of directing PKA phosphorylation to the bound substrate if the catalytic subunit must be released in order for phosphorylation to occur. Regardless of how close the catalytic subunits are positioned upon release, the CTP does not differ from that of randomly placed catalytic subunits. Once released, the CTP is equal to the expected CTP for catalytic subunits with access to the entire compartment that it is located in. Even for the very small volume of our simulation space, the CTPs for catalytic release are much slower than the rates for all tether models with the WT length (min 0.059s ±0.007 to max 0.087s ±0.011).

For some cell types, like the cardiac myocyte, AKAP directs phosphorylation to substrates that are distributed throughout the cytosol, thus the released catalytic subunit is not confined to a small compartment as in our LD simulations. In this case, the CTP for the substrate that the catalytic subunits were initially tethered to would decrease rapidly due to diffusion away from the initial location. Using the quasi-steady state Michaelis-Menten model of enzyme kinetics, we are able to generate an analytical prediction for the rate of substrate phosphorylation that could be expected given the volume of the compartment and presence of a single AKAP-PKA-substrate complex. While the assumptions implicit in the original derivation of Michaelis-Menten kinetics are not valid for our single molecule simulations, it has been demonstrated both theoretically and experimentally that this model is a good approximation of reaction velocity even for single enzyme kinetics [15, 16]. Using this method, we predicted the change in the CTP over time as the catalytic subunits diffuse away from their initial position (Fig 2B). The predicted reaction velocity was then converted to a catalytic rate (inverse of CTP) by dividing by the local concentration of the catalytic subunit. The analytical predictions of CTP agree reasonably well with the LD simulation results. The change in concentration of catalytic subunit over time was determined for a spherical volume with a radius equal to the mean squared displacement (MSD) of the catalytic subunit (Eq. 8).

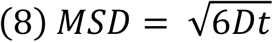

Once released, the catalytic subunit quickly diffuses away resulting in a drastic increase in the CTP (Fig 2B). The increase in CTP is such that after a few ms it would be unreasonable to expect that the catalytic subunit would return to its initial location and phosphorylate the AKAP bound substrate. Importantly, this demonstrates that outside of a small compartment like the dendrite of a neuron, release of the catalytic subunits from the AKAP-complex would be even less effective at achieving phosphorylation of complex bound substrates than for our small volume LD simulations.

### AKAP-PKA complexes with shorter more flexible tethers display faster characteristic times of enzyme-substrate binding and spend more time in the bound state on average

Phosphorylation of substrates by PKA occurs in a multi-step process involving binding of both ATP and substrate prior to catalysis. Due to the high cellular concentration of ATP, it is reasonable to model this process in two steps: 1) binding of PKA and substrate, 2) phosphorylation of substrate. Given that the catalytic rate of PKA is unlikely to be altered by the structure of the tether, we hypothesized the any changes in the apparent rate of catalysis would be driven by altered binding of the catalytic subunit with its substrate.

We first examined the characteristic time of binding (CTB), defined as the time from simulation start to the first binding event (Fig 3A). For this set of simulations, only the forward binding reaction was included (Fig 1C). CTBs varied from a minimum of 0.445ms ±0.078 to a maximum of 3.812ms ± 0.563. In general, shorter and more flexible tether lengths displayed faster CTBs. The effect of varying tether length is most dramatic for the least flexible model variations, while there is virtually no difference in the CTB with change in length for the most flexible model variations. Similar trends are seen with changes in flexibility at the extremes of the model variations. Changes in tether flexibility affect CTB least for short tether lengths and most for longer tether lengths.

**Figure 3.**
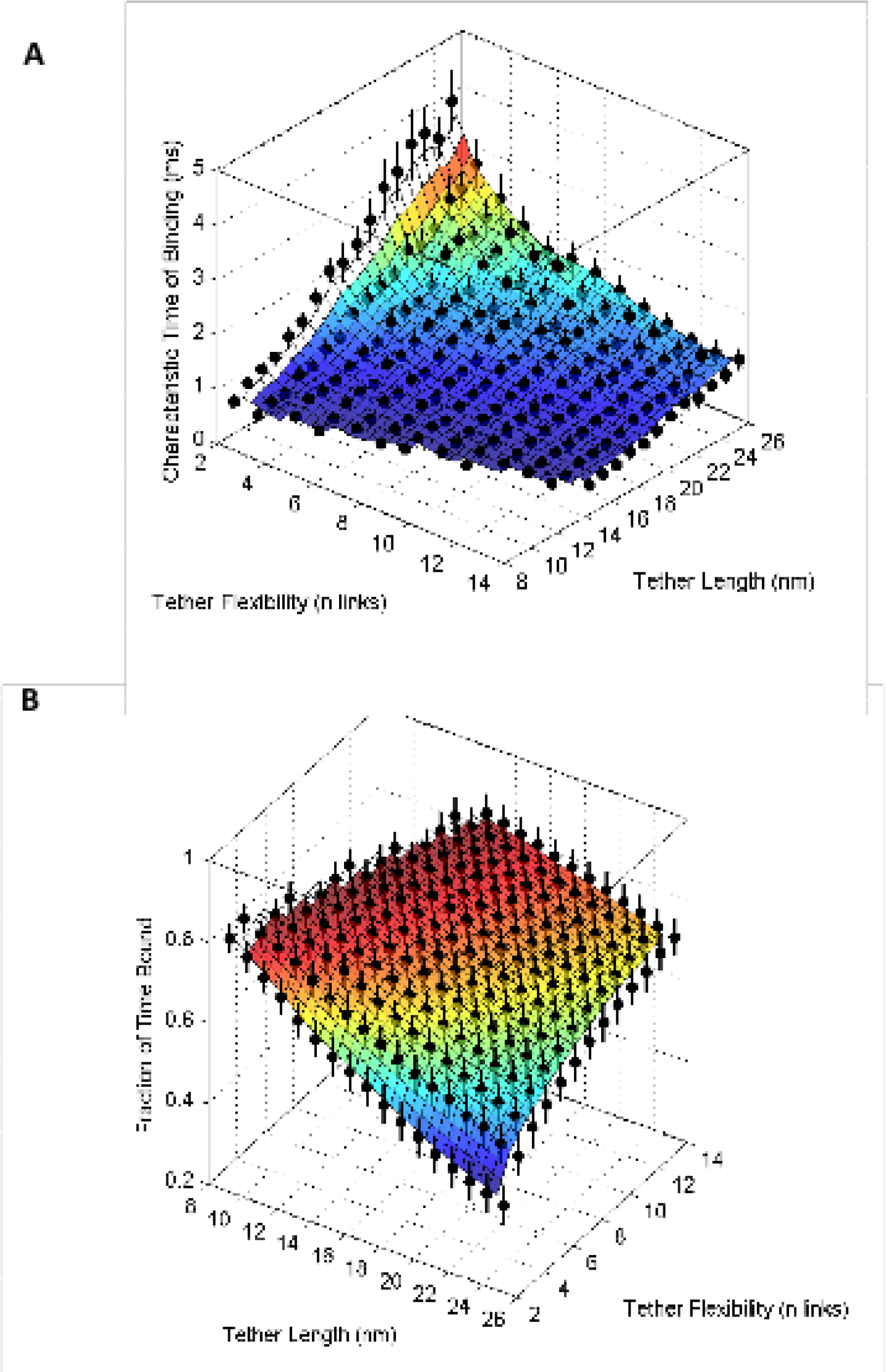
Tether length and flexibility modulate binding of the catalytic subunit to substrate. A) Characteristic times of binding with varied tether length and flexibility. Simulations were run for only the binding reaction of catalytic and substrate (no unbinding). Each data point represents the average time from simulation start to binding of catalytic and substrate for 10 sets of 50 simulations ±1SD. B) Average fraction of time that the catalytic subunit and substrate spend in the bound state. Binding and unbinding of the catalytic subunit with substrate were simulated over the course of 1s. Each data point represents the average fraction of time(s) ±1SD for 500 simulation runs.

Since the catalytic rate of PKA (21s^-1^) is much slower than k_r_ for binding of catalytic to substrate (350s^-1^), binding and unbinding of the catalytic subunit and substrate is expected to occur multiple times prior to phosphorylation on average. Model variations allowing catalytic and substrate to exist in a bound state for a greater fraction of total simulation time were hypothesized to have an increased probability of substrate phosphorylation and hence faster characteristic times of phosphorylation (CTP). We determined the average fraction of time spent in the bound state (FTB) by simulating binding and unbinding of catalytic subunit and substrate without allowing substrate phosphorylation (Fig 1C). The average FTB ranged from 0.415 ±0.05 to 0.847 ±0.043 with variations following the same trends as for characteristic times of binding (Fig 3B). Since the k_f_ and k_r_ remain unchanged for all model variations, the differences in the FTB must result from a change in the apparent forward rate of binding.

### Increases in apparent k_f_ of binding and average time spent in the bound state translate to decreases in characteristic times of phosphorylation

The variation in the CTBs and FTBs suggested that changes in the tether length and flexibility were likely to affect substrate phosphorylation. Model variations displaying an increased FTB were expected to display shorter CTPs because this should increase the probability of phosphorylation. We determined the CTP by simulating binding and unbinding of the catalytic subunit to substrate, and irreversible substrate phosphorylation (Fig 1C). CTPs varied from a minimum of 0.054s ±0.004 to a maximum of 0.119s ±0.012 (Fig 4A). As expected, the general trend was that models with shorter and more flexible tethers to displayed the fastest CTPs.

**Figure 4.**
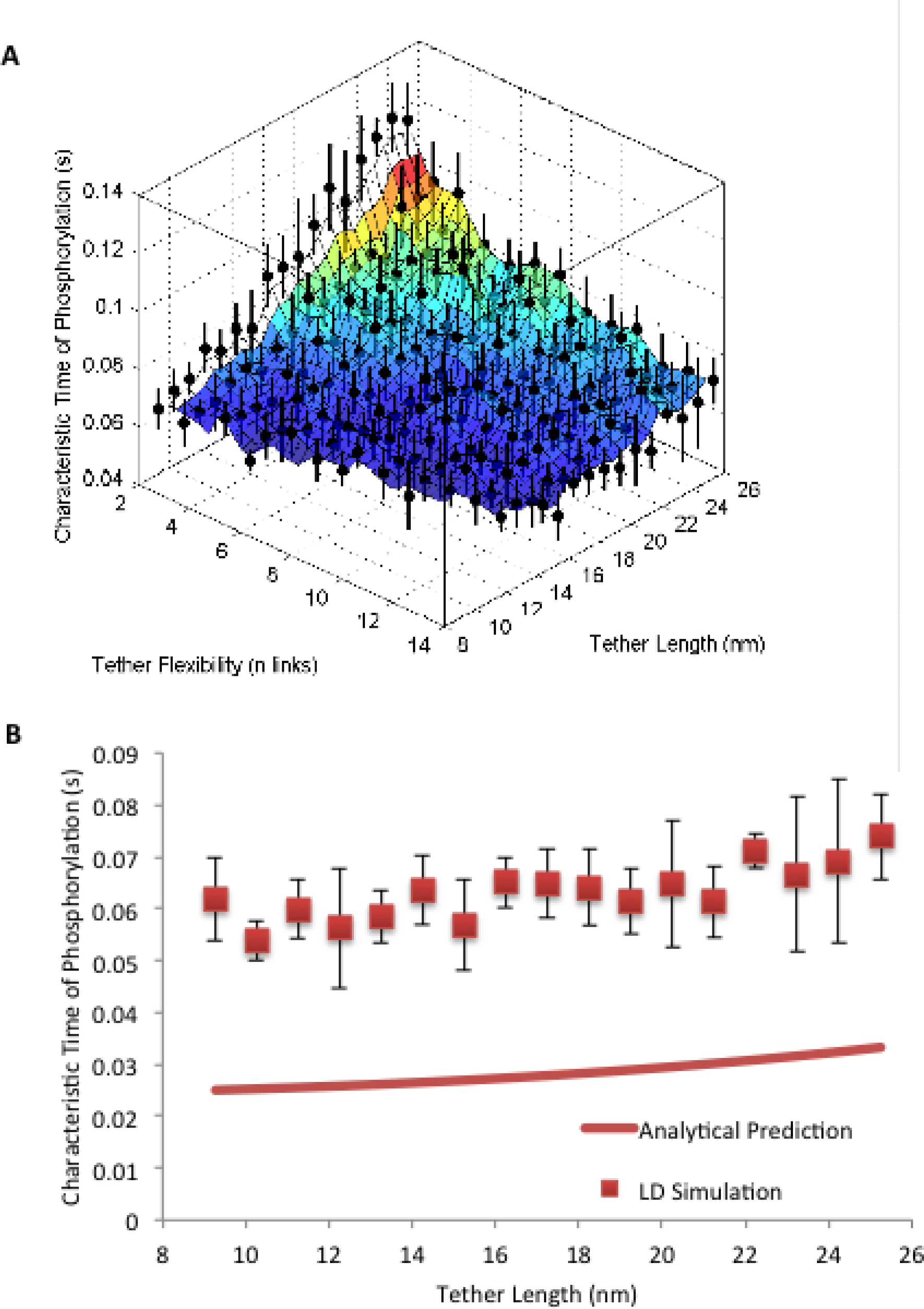
Tether length and flexibility modulate the rate substrate phosphorylation. A) Characteristic times of phosphorylation with varied tether length and flexibility. Simulations included binding and unbinding of the catalytic subunit with substrate as well as irreversible substrate phosphorylation. Each data point represents the average time from simulation start phosphorylation of substrate for 10 sets of 50 simulations ±1SD. B) Comparison of characteristic times of phosphorylation for the most flexible model variations at each length to characteristic times of phosphorylation predicted by Michaelis-Menten kinetics with adjusted concentrations of enzyme and substrate. A second set of LD simulations of the most flexible tether variations with only one catalytic subunit was run for each length for comparison to the analytical predictions.

In order to verify that the CTPs predicted from the LD simulations are realistic, we used the quasi-steady state Michaelis-Menten model of enzyme kinetics to generate an analytical prediction for comparison. While the assumptions implicit in the original derivation of Michaelis-Menten kinetics are not valid for our single molecule simulations, it has been demonstrated both theoretically and experimentally that this model is a good approximation of reaction velocity even for single enzyme kinetics [15, 16]. In order to determine the expected CTP using the Michaelis-Menten model, we adjusted the concentrations of the catalytic subunit and substrate by considering their local concentrations to be the number of molecules within a spherical volume having a radius equal to the length of the tether. The predicted reaction velocity was then converted to a catalytic rate (inverse of CTP) by dividing by the local concentration of the catalytic subunit.

The analytical predictions of CTP agree reasonably well with the LD simulation results for models with the most flexible tethers for each length (Fig 4B). The characteristic times of phosphorylation predicted by Michaelis-Menten kinetics are roughly twice as fast as those predicted by LD simulation. This deviation from the analytical prediction is a result of anchoring of the substrate to the membrane, which prevents its free diffusion within the same volume occupied by the catalytic subunits thereby reducing the CTP.

## Discussion

The importance of AKAPs in compartmentalization of cAMP-PKA signaling has been well demonstrated [18-20]. It is now widely accepted that AKAPs not only tether PKA to a particular substrate, they also co-localize with other signal effectors like adenylate cyclase and phosphodiesterase, thereby acting to coordinate a signaling microdomain [8]. Even though great progress has been made in our understanding of the spatial coordination of signaling components by AKAPs, how these AKAP complexes operate at the single molecule level remains poorly understood. While AKAPs are named for their ability to direct PKA phosphorylation to a particular substrate, we still do not have a molecular mechanism to explain how this occurs. This phenomenon is further complicated by years of evidence suggesting that activation of PKA requires the release of a freely diffusible catalytic subunit. However, recent evidence suggests that in the cell, the catalytic subunit is capable of phosphorylating AKAP complex bound substrates without being release [1, 2, 11]. It has been further demonstrated that structure of the PKA regulatory subunit tether is an important determinant of substrate phosphorylation [2].

We used Langevin Dynamics simulation of the AKAP-PKA-substrate complex to first identify whether or not tethering of PKA to its substrate in the AKAP-PKA-substrate complex was an effective method to ensure phosphorylation of the substrate if the catalytic subunit is released upon PKA activation. Next, we explored the contribution of the structure of the PKA regulatory subunit tether to substrate phosphorylation with the hope of gaining additional mechanistic insight into the function of the AKAP-PKA complex. Our work first demonstrates that positioning of the catalytic subunits within the space available to the WT tether length does not result in CTPs that are significantly different from those positioned randomly throughout the simulation space, indicating that AKAPs do not effectively direct PKA phosphorylation to AKAP bound substrates if the catalytic subunits are released (Fig 2A). We further demonstrate that in a large compartment like the cytosol, release of the catalytic subunit from the AKAP complex would lead to its quick diffusion away from the anchored substrate without phosphorylation (Fig 2B). The results of our simulations suggest that in the cell the catalytic subunit is likely not released, as this would lead to a significant decrease in the efficiency of phosphorylation of AKAP complex bound substrates. It is important to note, that in our simulations the substrate was almost never phosphorylated upon its first binding event with the catalytic subunit. This is a simple matter of reaction kinetics, given that the dissociation rate (350s^-1^) is roughly 17 times faster than the rate of catalysis (21s^-1^). This presents a strong argument for retention of the catalytic subunit within the AKAP-PKA-substrate complex, thus allowing for repeated interaction of catalytic subunit and substrate that will increase the probability of phosphorylation.

Next, we demonstrate that modulation of the tether length and flexibility changes the CTB between the catalytic subunit and substrate, with shorter and more flexible tethers tending to have faster CTBs (Fig 3A). The change in CTB, resulting only from alterations in tether structure, modulates the fraction of time that catalytic subunit and substrate spend in a bound state (Fig 3B). This translates into changes in the CTP, with models having higher FTBs showing decreased CTPs (Fig 4A).

The decreased CTB and increased FTB for shorter tether lengths reflect an increased apparent forward rate of binding. Since the rate constants (k_f_ and k_r_) remain the same for all model variations, the changes in the apparent forward rate indicate that the structure of the tether influences the probability of interaction of the catalytic subunit with its substrate. As the regulatory subunit tether is shortened, the space available to the catalytic subunit decreases, thereby increasing the probability that it will encounter and bind to the substrate as long as the substrate remains accessible. The increased probability of interaction has previously been interpreted as resulting from an increase in the local concentration of the catalytic subunit. While this interpretation allows us to easily predict the CTP via the Michaelis-Menten model, the changes in CTB, FTB, and CTP result from differences in structure, not concentration, and thus are more correctly thought of as changes in apparent forward rate constant. The flexibility of the tether likely modulates accessibility of the space within the spherical volume defined by the tether length. For lengths that allow access to substrate in the most extended conformation, flexibility has little impact on CTB or FTB. With longer tethers that must deform in order to reach the substrate, changes in flexibility have much more impact on the CTB and FTB (Fig 3). Although differences in the regulatory subunit tether length are only expected with interspecies variation, positioning of substrates within the AKAP complex does vary and would affect rates of interaction similarly. One potential caveat to bear in mind is our inability to account for rotational conformation. In reality, phosphorylation of substrates by the catalytic subunit would require precise alignment of the catalytic cleft with the substrate. It may be possible that a certain degree of tether stiffness could actually facilitate phosphorylation by maintaining proper alignment of the catalytic subunit and substrate.

Changes in the CTP with tether length and flexibility are most likely driven by changes in the apparent forward rate of binding. Although the idea of a local concentration offers a simple method for analytical prediction of the CTP, the actual concentration of the catalytic subunit and substrate do not change. Considering the changes CTB and FTB as resulting from changes in apparent k_f_ rather than concentration offers interesting insight into the changes in CTP and, more generally, the function of AKAPs. Increasing the k_f_ of the binding reaction results in a decrease in the Michaelis constant (Methods Eq. 7), which translates into an increase in the velocity of the phosphorylation reaction according the Michaelis-Menten model (Methods Eq. 6). Interestingly, such change in the Km also suggests that the efficiency of phosphorylation is improved according to the following equation (Eq. 9), where E is catalytic efficiency:

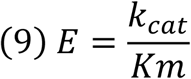

This interpretation of the change in CTP resulting from changes in tether structure is a previously unconsidered, but incredibly important advancement in our understanding of the function of AKAPs. It suggests that AKAPs don’t simply increase the speed or magnitude of phosphorylation by increasing local concentration of PKA, rather AKAPs increase the efficiency of phosphorylation of the AKAP complex bound substrate.

The increase in efficiency of phosphorylation of AKAP bound substrates is best illustrated by comparing the CTPs of tether models with those from our models of catalytic release. For models involving catalytic release, the CTP is at minimum 2.5-fold slower than the slowest tether model. Given the small size of our simulation space and our evidence that the rate of substrate phosphorylation depends on the size of the compartment for free catalytic subunit, the increase CTP is likely to be much more dramatic for most AKAP-PKA signaling systems in the cell.

Our new hypothesis regarding the function of AKAPs also offers some interesting interpretations of old data. While the definition of local concentration of PKA in the previous hypothesis is somewhat ambiguous, we will define it as concentration of PKA within the compartment of the AKAP-PKA complex. Experiments in which AKAP-substrate binding has been disrupted demonstrate a reduction in phosphorylation of the intended substrates [19, 21]. While this was attributed to de-localization of PKA and hence, a decrease in local concentration, our evidence indicates that such delocalization would also occur upon activation of PKA if the catalytic subunit is released from the complex in order for phosphorylation to occur. The effect of tethering the catalytic subunit to its target is even better demonstrated through the use of the FRET sensor AKAR (A Kinase Activity Reporter 2), which measures PKA phosphorylation real time in cells [22]. Addition of the PKA binding site to the AKAR2 construct resulted in FRET signal changes following stimulation that were both faster and of higher magnitude when compared to AKAR2 alone [11]. Since AKAR2 with or without the addition of the PKA binding site is localized only to the cytosol, it is not reasonable to assert that there is difference in the local concentration of PKA. The sensor is located in the same compartment in both cases and we are not forcing redistribution of PKA to a larger space by preventing AKAP localization to a particular substrate. Our hypothesis that AKAPs increase the efficiency of AKAP complex bound substrate phosphorylation fits very well with this data. Our simulation data suggests that the increased rate of phosphorylation, and likely the increased magnitude, is what should be expected. Interestingly, we find that the kinetics of phosphorylation for released catalytic subunit are too slow to mediate some signal processes known to require PKA. The time scale of PKA mediated effects on Na^2+^ channels occur on the order of milliseconds, which according to our LD simulations occurs too quickly to be mediated by the catalytic subunit if released from the complex [23].

Computational analysis of the AKAP-PKA complex using our novel software, *SpringSaLaD*, has led to numerous exciting findings. First, we present evidence that release of the catalytic subunit would likely abrogate the effect of tethering PKA to a particular substrate, supporting the recent findings of *Smith et al* [1, 2]. This evidence suggests that for efficient substrate phosphorylation to occur, the catalytic subunit cannot be released. Given the low probability of substrate phosphorylation upon first interaction with catalytic subunit, it is unreasonable to expect that a free catalytic subunit would efficiently phosphorylate the substrate it was initially tethered to. Second, we offer a mechanism for the changes in phosphorylation rate seen with alteration in the structure in regulatory subunit tether. This change in the mechanism of phosphorylation also provides us with a partial molecular mechanism that explains how AKAPs might direct PKA phosphorylation to AKAP complex bound substrates. Finally, we propose an improvement to the hypothesis that AKAPs increase the speed and magnitude of substrate phosphorylation by increasing local concentration. We hypothesize that AKAPs act to increase the efficiency of PKA phosphorylation and suggest that “local concentration” should be thought of as the amount of enzyme and substrate within a volume defined by the length of the regulatory subunit tether, rather than within a local region or compartment within the cell. This revision of the AKAP hypothesis represents a critically important step forward in our understanding of PKA signaling.

## Author Contributions

MR carried out the simulations, analyzed the data, and wrote the manuscript, PJM contributed to simulation software configuration, MR, KLD-K, IIM designed the experiments, interpreted the data, and edited the manuscript.

## Acknowledgements

This work was supported by NIH grant P41-GM103313 (IIM) and an American Heart Association Grant 12GRNT1216008 (KLD-K).

